# Agents seeking long-term access to the wisdom of the crowd reduce immediate decision-making accuracy

**DOI:** 10.1101/2023.09.16.558066

**Authors:** Richard P. Mann

## Abstract

Living in groups offers social animals the significant advantage of accessing collective wisdom and enhanced information processing, enabling more accurate decisions related to foraging, navigation, and habitat selection. Preserving group membership is crucial for sustaining access to collective wisdom, incentivising animals to prioritise group cohesion. However, when individuals encounter divergent information about the quality of various options this can create a conflict between pursuing immediate rewards and the maintenance of group membership to improve access to future payoffs. In this study I show that rational agents who seek to maximise long-term rewards will be more inclined to follow the decisions of their peers than those with short-term horizons. In doing so, they necessarily make less-rewarding decisions in the short-term, which manifests in a lower individual accuracy when choosing the better of two options. Further, I demonstrate that intuitions about collective wisdom can be misleading in groups of agents who prioritise long-term rewards, with disagreement being a better signal for the accuracy of collective choices than consensus. These results demonstrate that observed patterns of sociality should be interpreted in the context of the life history of an individual and its peers, rather than through the lens of an isolated decision.

## Introduction

Group living confers a multitude of advantages upon social animals [1, 2]. Among these, the ability to leverage collective intelligence and social information stands out as a significant benefit, serving to enhance the quality of decision-making processes [3, 4, 5]. For animals living in groups, the behaviour of other individuals is an important source of information about which actions are likely to lead to rewards such as food, shelter and safety. A decision by one animal to move towards a specific location, or to forage at a specific time, provides a credible signal about its knowledge of likely food abundance and risk in that time and place [6, 7]. A second individual in the same group can combine that signal with its own information to make better choices than it could do alone: optimising food intake, minimising predation risk, and thus maximising its fitness. As such, even in groups of unrelated individuals, information sharing is an important benefit of group living that provides a powerful incentive to remain together.

The use of social information in decision-making can be understood as an adaptive behaviour that maximises fitness by enabling an animal to choose behaviours with the highest expected reward, whether through an explicitly cognitive process or via heuristic ‘rules of interaction’ – ingrained, evolutionarily-selected tendencies to copy what others do under certain condition or consistent weightings of social and non-social information [8]. Following this logic, previous research has sought to derive these rules of behaviour from first principles, by considering what a rational agent would do given the social and non-social information available when faced with a decision between different options with uncertain rewards [9, 10, 11, 12, 13, 14, 15]. These studies have focused on optimising an agent’s expected reward in a single decision, while treating the group’s size and composition as independent variables, unrelated to the decision at hand. However, the decisions animals make are potentially drivers of group composition. An individual that chooses to move in a direction opposed to others may find itself isolated for some period of time, and thus faced with making further decisions alone without access to the benefits of collective wisdom. Even without a spatial context, disagreements can fracture groups in contexts where homophily is a factor in determining social connections [16, 17]. Since being in a group allows an agent to access beneficial social information, this provides an incentive to follow the decisions made by others even if an individual has reason to believe that higher immediate rewards are available elsewhere, introducing a trade-off that is ignored by normative mathematical theories of social decision-making that focus on isolated choices. This is in sharp contrast to other frameworks for modelling optimal decision-making under uncertainty, such as reinforcement learning, in which expected rewards in the present are weighed against potential rewards in the future, with a ‘discount rate’ specifying the relative importance of each. In this paper I develop a model of collective decision-making by rational agents that incorporates the link between individual decisions and group membership, and explore the consequences of this connection for group cohesion and the expected accuracy of individual and collective decisions when agents value future rewards more or less highly.

## Model

I first consider a simple model of two agents making repeated binary decisions. By considering just two agents this reduces the problem to its most fundamental constituents, as well as making mathematical treatment more tractable. Subsequently I show how the model can be extended to the case of larger groups.

In each decision there are two options, A and B, one of which is associated with a reward of unit value (in arbitrary units of utility). I define *x* as an indicator variable for the correct choice, such that *x* = 1 if A is correct, and *x* = *−*1 if B. Before observing any information, agents share a common, uninformative prior belief about the value of *x*:

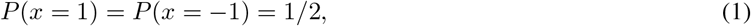

Each agent subsequently receives a private signal about the value of *x*, labelled as Δ_*i*_, centred on the true value of *x* and normally distributed with noise variance *ϵ*^2^:

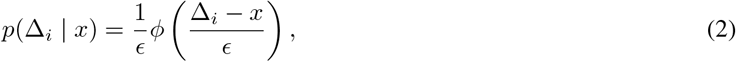

where *ϕ*(*·*) is the standard normal probability density function. Agents are assumed to receive signals of equal reliability (*ϵ* is the same for all agents), and this is equivalence is assumed to be common knowledge [18]. The two agents choose between A and B in a randomly-chosen order, and the first agent has only their private information on which to base their choice. Therefore, the first agent will choose A if and only if Δ_1_ *>* 0, since this implies that *P* (*x* = 1 | Δ_1_) *>* 1*/*2. The second agent can use the decision of the first to inform its own choice, by setting a threshold on its private information Δ^*∗*^, such that it will select option A if and only if Δ_2_ *> C*_1_Δ^*∗*^, where *C*_1_ indicates the decision made by the first agent (1 for A, *−*1 for B). For a given threshold value, one can derive the expected reward each agent can expect to receive in a given decision; that is, the proportion of decisions where the agent is expected to choose the correct option. For the purpose of calculation, we can assume without loss of generality that for a given decision option *A* is better, and with this assumption the expected reward 𝔼_*T*_ (*R*) is:

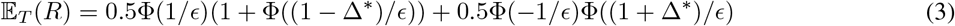

where Φ(*·*) is the standard cumulative normal probability function. Here the *T* subscript in the expectation indicates that this is the expected reward for agents choosing together, and the expected reward is averaged over both agents, since I assume each agent is equally likely to choose first or second. Maximising this expectation with respect to Δ^*∗*^ provides the optimal threshold value, which matches that derived in ref. [19].

The expected reward above can be compared to that for a single agent choosing alone. In this case no social information threshold is relevant, and the expected solo reward 𝔼_*S*_(*R*) is:

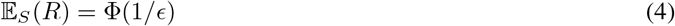

### Repeated decisions

I now consider the case where an agent will expect to make multiple decisions over time, and will seek to maximise its expected reward across the sequence of all of these. For the sake of simplicity and mathematical tractability, I assume that all decisions an agent will face are of equal difficulty (i.e. *ϵ* remains constant throughout) and have equal potential payoffs.

I introduce a new parameter *γ*, which denotes the probability that each decision will not be the last in this sequence, such that the expected number of future decisions at any time is 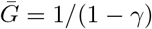. The mathematical analysis below is also the same if *γ* is treated as a discount rate, such that the agent treats future rewards as less valuable than immediate ones. In either case, the expected total reward is given by:

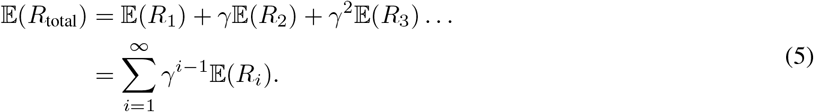

In the equation above the expectations do not have subscripts indicating whether the agents are choosing together or alone. In the key aspect of the model, I assume that the agents are initially together, and remain together for as long as they make identical decisions. When they make differing decisions they must henceforward make their decisions alone. Therefore we can expand the expectation above as:

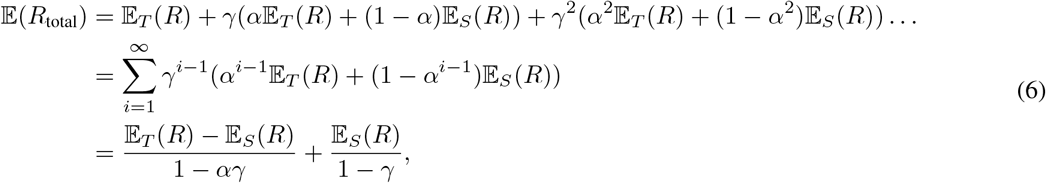

where *α* is the probability for the two agents to make the same decision and thus remain together:

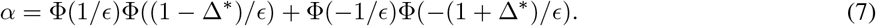

By maximising equation 6 with respect to Δ^*∗*^ we can identify the optimal value of this threshold that maximises the expected total reward for an agent over the full sequence of future decisions.

### Larger group sizes

In a group of *n* agents choosing between two options there are *n* + 1 possible aggregate outcomes, ranging from 0 to *n* agents selecting each option. Where the outcome of the current decision determines the group that an agent will find itself in for subsequent choices, that agent must evaluate how beneficial it will be to be in a group of 1, 2, … *n* at the next iteration of the game, and the probability of finding itself in that situation based on its current decision. Moreover, each agent can observe 2^*n*^ *−* 1 different configurations of social information, in the form of all possible sequences of 0 to *n −* 1 binary decisions, each of which can be associated with a different threshold value for their private information (though symmetry between sequences can reduce this number to 2^*n−*1^ *−* 1). To derive an optimal strategy one must therefore proceed in stages.

First we specify the expected lifetime reward of a group of one (where lifetime is taken to mean the series of games the agent will play):

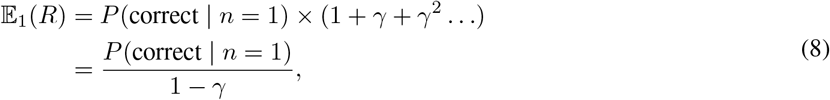

where *P* (correct | *n* = 1) denotes the probability of choosing correctly in a group of size 1. This expected reward is simply the probability of choosing the correct option in a single game when playing alone, multiplied by the expected number of games to be played. This can be rewritten as:

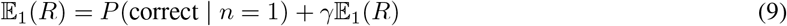

This form indicates that the expected reward is composed of the probability of choosing correctly in the current game, plus all the expected rewards that will accumulate if the series of games continues, with probability *γ*.

The optimal strategy for an agent alone is clear: it should seek to maximise the probability of making the correct choice in the first game, since the probability of the series of games continuing does not depend on the choice it makes. Thus, we can immediately specify that the critical value of 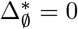 (where the subscript denotes the sequence of previous decisions the agent observes, and ∅ indicates the empty set), and thus

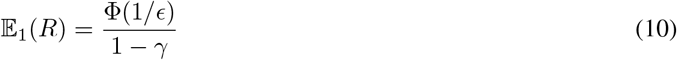

For agents in a group of two, the optimal strategy for each agent depends on which position it finds itself in for the current game and the sequence of previous decisions it observes. Denote as 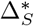 the critical value employed when observing a sequence *S*, and note that by symmetry 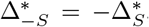, where *− S* indicates the sequence *S* with all decisions reversed. Then in a group of two there are only two critical values to determine: 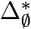 for the first agent, who observes no prior decisions, and 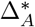 for the second agent after observing the first agent choose *A* (since 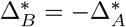). Again considering the symmetry of the game, it is clear that 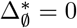 continues to apply for all group sizes. Therefore, it remains to determine 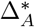. For the second agent, its expected total reward is given by:

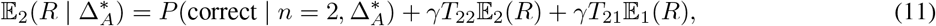

where *T*_22_ and *T*_21_ are elements of a transition matrix, indicating the probability that an agent in a group of two will subsequently find itself in a group of two or one at the next game, and are understood to depend on 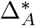. If the focal agent selects 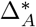 optimally it can assume that this value will be applied consistently in the future by other agents in the same situation, so *T*_22_ and *T*_21_ can be assumed to be constant in time. Assuming without loss of generality that the true correct choice is *A* (*x* = 1), and considering all sequences of events and potential values of private information the two agents may observe, the key terms in this equation are given as:

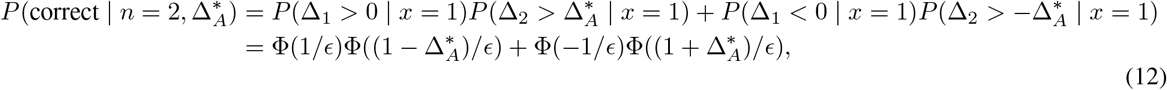

and

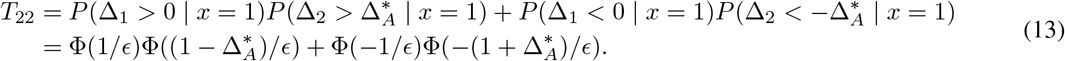

Thus we recover the previously derived optimal strategy for two agents, with *α* = *T*_22_ and 1 *− α* = *T*_21_.

With three or more agents, deriving the optimal strategy is more complex, since the decision an agent makes may influence many subsequent decisions by other agents, and thus the probability of remaining in different group sizes for subsequent decisions. To obtain the optimal strategy in a group of size *n*, we assume that the expected reward of a randomly chosen agent in groups of size 1, …, *n −* 1 are known; this assumption means that we must first identify the optimal strategy for these smaller groups. We then initialise the critical value associated with all observable sequences to zero: 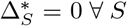. Based on these initial values, we can calculate the probability of generating any sequence of decisions given a true value of *x* = 1 as:

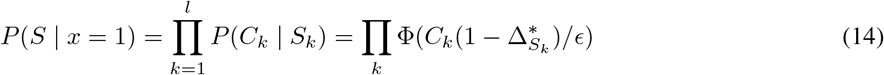

where *C*_*k*_ is the *k*th element of *S, S* = {*C*_1_, … *C*_*l*_} and *S*_*k*_ = {*C*_1_, … *C*_*k−*1_} is the subset of *S* composed of the first *k −* 1 decisions. By evaluating all the probability of all sequences of length *l* = *n*, we can determine the expected proportion of times that an agent that observes sequence *Z* will choose correctly and the expected number of other agents it will agree with (which forms the group size in the next game), and thus the transition probabilities *T*_*ij*_. We then consider varying the value of this focal critical value, 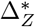 so as to maximise the expected lifetime reward for this agent (while maintaining the symmetrical condition that 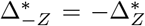). By successively optimising the expected lifetime reward for each observable sequence, we arrive at a new complete set of values 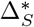. Repeating this process until convergence to stable values, we can determine the optimal strategy for all agents observing any sequence of previous decisions.

Having obtained the optimal values of 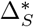 for all sequences *S*, we can finally determine the probability that a focal agent, observing sequence *S*, will choose option A, conditioned on the value of *x* that indicates the true correct option:

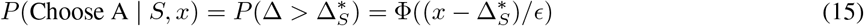

## Results

### The effect of future-weighting on rational decision rules

To illustrate the effect of changing the value of the future weighting (*γ*) I consider the probability for a focal rational agent to choose option A, conditioned on the number of previous agents choosing either A (*n*_*A*_) or B (*n*_*B*_). This is a compression of the actual decision rule the agents obey, since it omits details of the precise sequence of choices that the agent can observe. It should therefore be stressed that agents respond to the full, ordered sequence of previous choices and the choice to illustrate this response in a lower-dimensional form is motivated by two considerations. First, a pragmatic choice that provides ease of visualisation. Second, for comparison to both previous theoretical work that utilises similar characterisation (e.g. ref. [12]), and to empirical work in which the ordering of decisions is often omitted (e.g. refs. [20, 21])

Without loss of generality, I assume arbitrarily that A is the correct choice, such that *x* = 1. The same number of agents choosing each option can be obtained from multiple different sequences of decisions, for example *n*_*A*_ = 2, *n*_*B*_ = 1 is consistent with sequences AAB, ABA and BAA; I therefore average over these sequences according to the probability of their occurrence to obtain the probability conditioned on *n*_*A*_, *n*_*B*_:

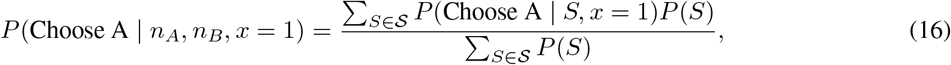

where the summation is over all sequences *S* in the set of sequences 𝒮 consistent with the observed values of *n*_*A*_, *n*_*B*_.

Figure 1 visualises this conditional probability for three values of the future-weighting: *γ* = 0 (panel A), *γ* = 0.9 (panel B) and *γ* = 0.99 (panel C), in a group of 15 agents. These show that as *γ* increases the focal agent becomes increasingly likely to follow the majority of previous decisions. This is reflected in the predicted collective outcome - the probability for a given total number of agents to select option A (*N*_*A*_), shown in panels D-F. These show an increasing tendency to consensus as *γ* increases, such that when *γ* = 0.99 (representing an expectation of 100 future games) the agents are overwhelmingly likely to all choose the same option (whichever choice is made by the first agent). Notably, both the probability of all agents choosing correctly and the probability of all agents choosing incorrectly increase as *γ* increases.

**Figure 1.**
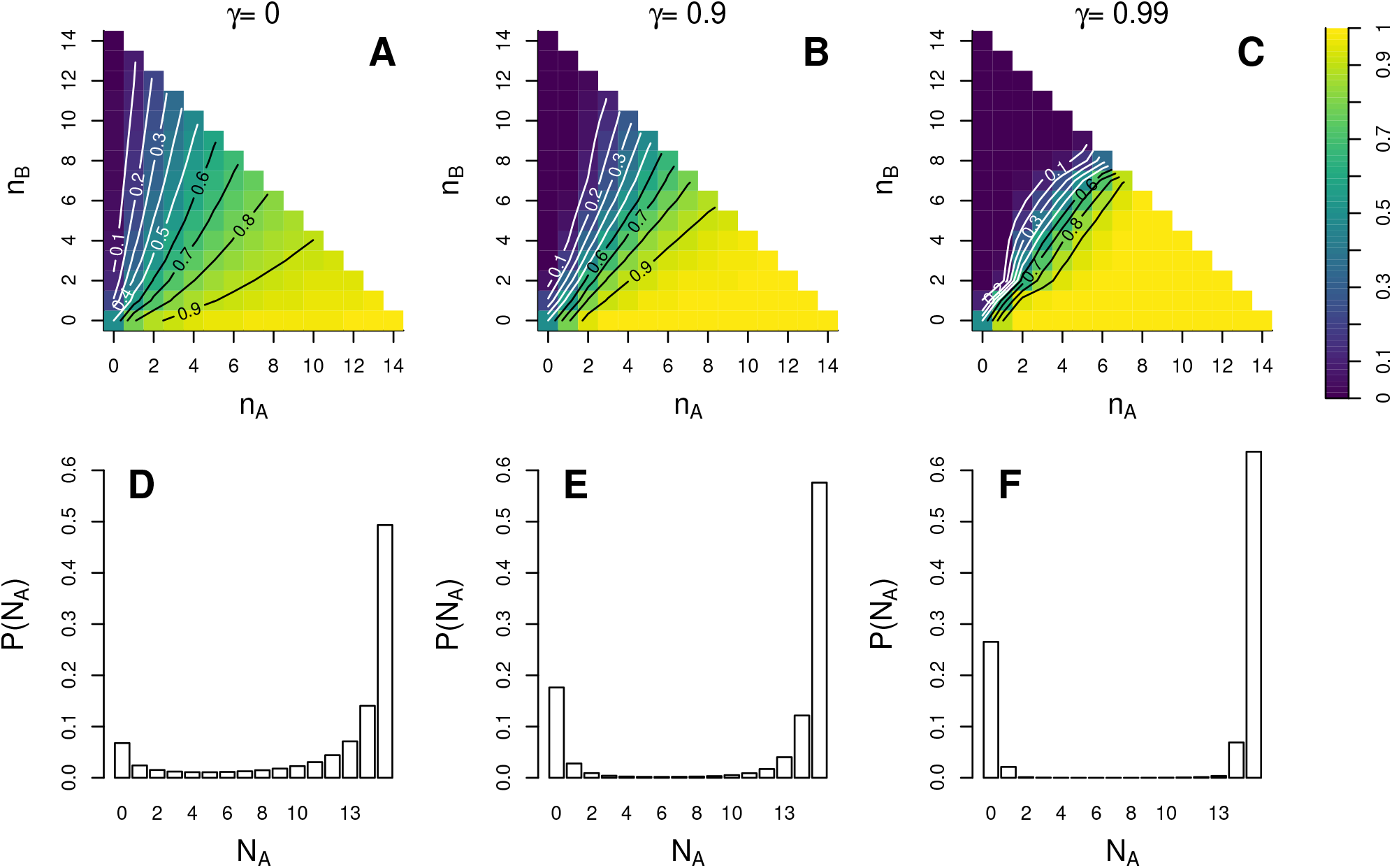
Characterising the rational strategy for agents in larger groups. Each column shows results for a different value of *γ*, with the environmental noise set to give a solo accuracy of Φ(1*/ϵ*) = 2*/*3. Top row: the probability for a rational agent to choose the correct option A (*x* = 1), conditioned on the number of previous decisions for each option (*n*_*A*_, *n*_*B*_), with future weightings of (A) *γ* = 0; (B) *γ* = 0.9; (C) *γ* = 0.99, corresponding to 1, 10 or 100 expected future games respectively. When agents prioritise immediate rewards, the social response is characterised by a sensitivity to the precise values of *n*_*A*_, *n*_*B*_; as the future weighting *γ* increases, rational agents become increasingly likely to follow the majority decision, with little sensitivity to the precise size of that majority. Bottom row: the collective outcome of agents employing the rational strategy, the probability that a total of *N*_*A*_ agents choose the correct option A. When agents prioritise immediate rewards (D), consensus decisions are likely, but the group often splits. As *γ* increases (E, F), consensus decisions become much more likely, and in particular the probability that the whole group chooses incorrectly increases strongly.

### Effect of future-weighting on individual accuracy

Agents that prioritise remaining with others over immediate rewards necessarily become less likely to make correct decisions in the present. To explore this effect I determined the optimal strategy for individuals in groups of one to ten agents as a function of the expected number of future games (1*/*(1 *− γ*)), and from this calculated how often a randomly chosen agent within the group will choose the correct option. These results, shown in Figure 2, show as expected that individual accuracy in the present is maximised when agents do not consider the future (*γ* = 0). As the expected number of future games to be played increases, agents’ average accuracy in a single game rapidly falls, approaching the accuracy for a single agent choosing alone (black line). However, for any value of *γ* it remains the case that larger groups exhibit greater average accuracy than smaller groups, illustrating the incentive for agents to remain in these larger groups to increase their own accuracy in future decisions.

**Figure 2.**
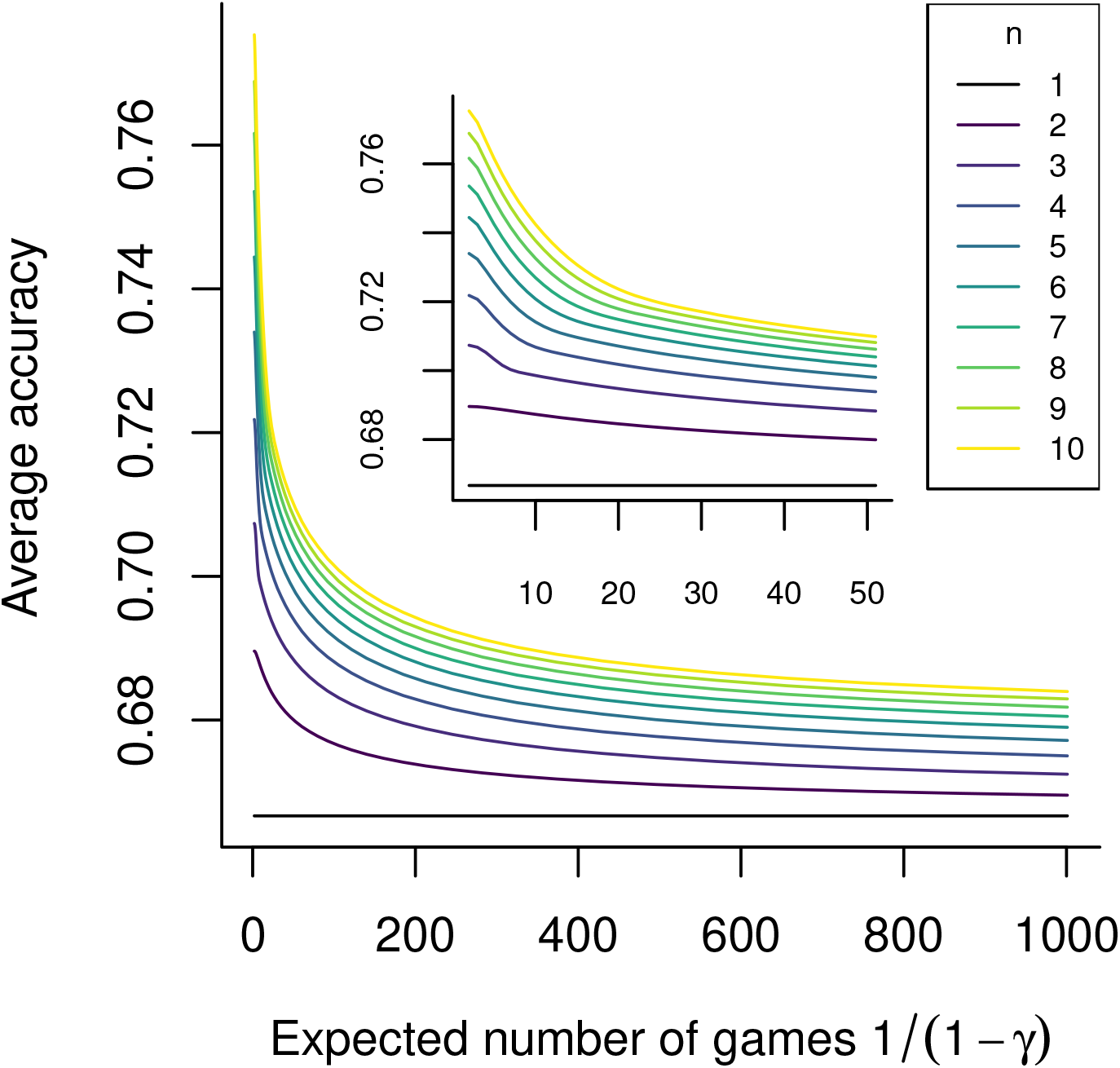
The average accuracy of individual decisions as a function of group size and future-weighting. Each curve shows the proportion of times a randomly selected agent in a group will make the correct choice in a single decision when agents adopt the optimal strategy, as a function of the expected number of future games 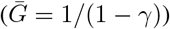for a specific group size; the inset shows the detail of these curves for values of 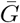 from 1 to 50. The environmental noise is set to give a solo accuracy of Φ(1*/ϵ*) = 2*/*3. For every group size, average accuracy is greatest when the expected number of future games is 0, since agents can prioritise immediate accuracy. As the expected number of games increases the average accuracy falls for all group sizes, as agents prioritise remaining with their peers over immediate accuracy. For any future weighting, larger groups give greater average accuracy, illustrating the incentive to remain in a larger group for future decisions.

It should be reiterated for clarity that although accuracy in a single game declines when agents anticipate playing many future games, their strategy is optimal by design and results in greater accuracy over the long term if those anticipated games come to pass. The loss of accuracy in present is compensated by an increased probability of remaining in a larger group for future games, which allows them to be more accurate in these later games because of the positive effect of group size. Thus the results shown in Figure 2 do not indicate that a high future weighting (*γ*) is detrimental, but only that it reduces the accuracy in the present as a necessary sacrifice to obtain greater expected rewards in the future.

### Collective accuracy in groups prioritising cohesion

Since the accuracy of individual decisions is reduced by agents prioritising continued group membership, this suggests that such groups may be less reliable as a source of ‘collective wisdom’ to an outside observer. The Condorcet Jury Theorem (CJT) states that if agents individually make independent choices with some accuracy greater than 0.5, the probability that the majority of the group chooses correctly rapidly approaches one. However, the assumption of independent decision-making in the CJT is already violated when rational agents choose sequentially, and further biases towards consensus tend to further decrease the accuracy of the majority decision [19]. To examine how the bias towards consensus induced by future-weighting affects collective wisdom, I calculated the probability of all decision sequences in a group of 15 agents for cases where the correct option was A (*x* = 1) or B (*x* = *−*1), varying the future-weighting in the form of the expected number of future games (1*/*(1 *− γ*)). I then evaluated the probability that A was the correct choice, conditioned on observing the total number of agents who selected that option, assuming that both options are equally likely *a priori*:

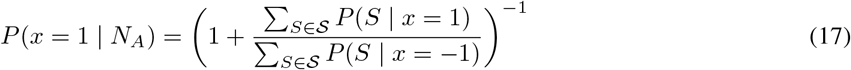

where the summations are over all sequences of decisions in the set *S* consistent with *N*_*A*_ total agents choosing A. It is worth reiterating at this point that the agents themselves are assumed to observe and respond to the full sequence of previous choices; the calculation above considers how much information the collective outcome of this process provides to an external observer who only observes the aggregate number of decisions for A and B.

As shown in Figure 3A, when only one game is considered (*γ* = 0), the probability that A is the correct choice, as evaluated by an outside observer, monotonically increases with the number of agents making that choice (black line). This under-performs relative to the information provided by agents choosing independently as per the CJT (grey line), but each additional agent choosing A continues to add weight to the belief that A is the correct choice. However, as the future-weighting increases this monotonic relationship breaks down. When many future games are considered by the agents the most informative outcomes are those in which the agents make differing choices. For example, when the expected number of future games is 100 (green line, *γ* = 0.99), the greatest certainty about the true correct choice is obtained when *N*_*A*_ = 6 or *N*_*A*_ = 9, i.e. a small majority is more trustworthy than a consensus. This is because a consensus decision often represents agents with relatively weak personal information strongly following those who choose before themselves, resulting in an information cascade that simply amplifies the decision made by the first agent. Conversely, small majorities (which occur rarely, because of the strong social response), represent occasions when later agents had sufficiently strong personal information to overcome incorrect choices made by the earliest decision-makers. Such strong environmental signals in favour of one option are much more likely when that option is truly correct, and so small majorities are much informative when strong signals are required to overcome previous social information. Very small majorities (e.g. *N*_*A*_ = 8, *N*_*B*_ = 7) can even be more informative under high values of *γ* than the same majority achieved by independent decision makers because of this effect, although such outcomes are very rare (see Figure 1E-F).

**Figure 3.**
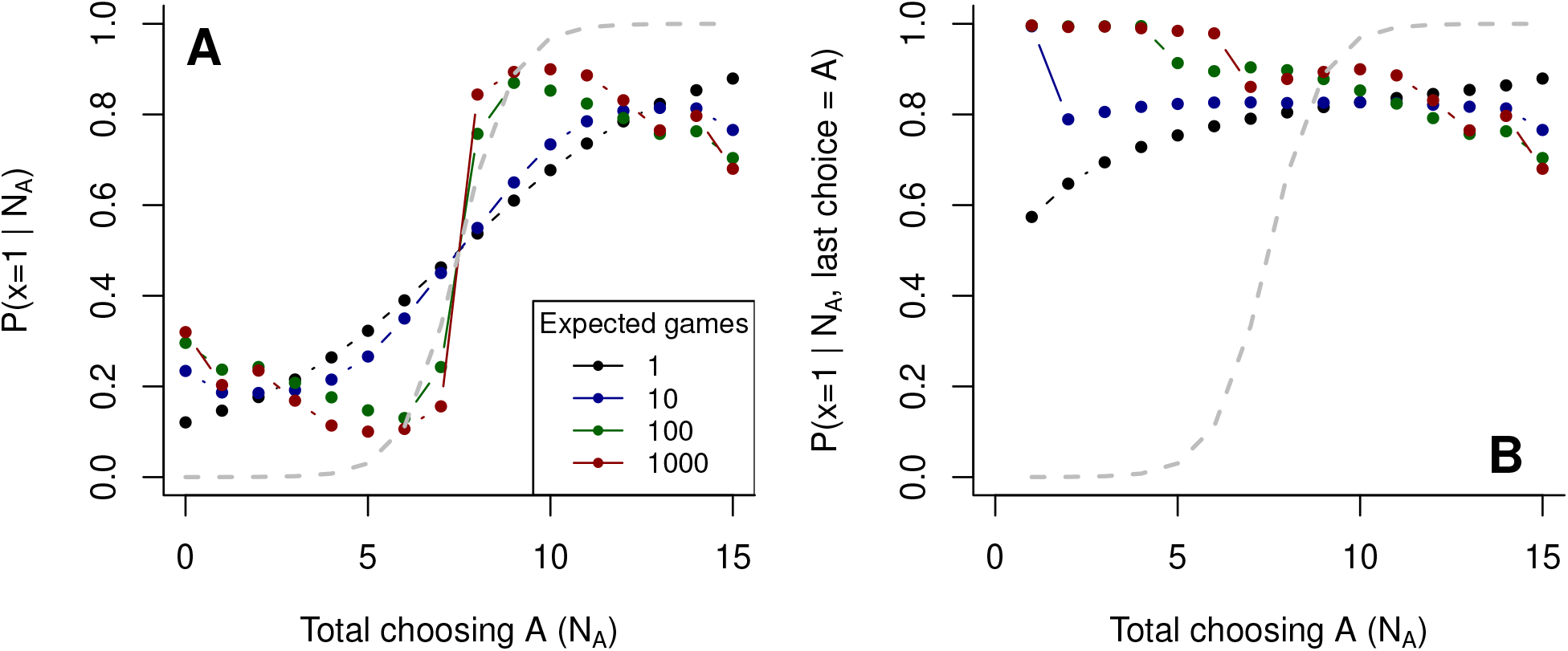
Information provided to an external observer from collective outcomes in groups of differing future weighting, with environmental noise set to give a solo accuracy of Φ(1*/ϵ*) = 2*/*3. (A) Each line represents the probability that option A is the correct choice, conditioned on the number of agents choosing A (*N*_*A*_) in a group of 15 agents. The grey line indicates the probability given by agents choosing independently, corresponding to the predictions of the Condorcet Jury Theorem (CJT). When agents choose independently the majority are highly likely to be correct, and larger majorities are more trustworthy. Each further line indicates results for a group of non-independent agents using the optimal strategy for different numbers of expected future games. When playing a single game (black line) the majority is less trustworthy than under the CJT, but larger majorities continue to be more trustworthy than smaller ones. As the expected number of future games increases, consensus decisions become increasingly untrustworthy, as they typically result from information cascades; for high values of *γ* the most trustworthy outcomes are (rare) small majorities, indicating that early, incorrect decisions have been overcome by more reliable private information among later agents. (B) Information provided by collective outcomes conditioned on the most recent decision being for option A. In all cases the probability that *x* = 1 is greater than 0.5 across all values of *N*_*A*_, indicating the high information content of the most recent decision. As the expected number of games becomes large the most informative outcomes are those where the final decision disagrees with the previous majority.

The results shown in Figure 3A show the information available to an external observer who is only able to observe the aggregate number of individuals choosing each option. However, in sequential decisions the order in which decisions were made can be highly informative. In particular, the most recent decision is often of greater importance than all previous decisions taken together (cf. ref. [12]), incorporating knowledge of this most-recent decision can provide an observer with essentially the same information as the full sequence of decisions [14]. To illustrate this effect, Figure 3B shows the information provided by collective outcomes conditioned on the final decision that was made. For the case where agents expect to play one game (*γ* = 0), the probability that the correct choice is A is always greater than 0.5 regardless of the actual number of agents choosing A, but increases with *N*_*A*_. However, when agents consider many future games the most informative outcomes are those where the final decision is A in spite of a large majority of previous agents choosing B. In these cases the final agent must have exceptionally strong private information in favour of A to overcome the previous social information. However, from the perspective of an external observer there is no ‘free lunch’ – these most informative outcomes are also very unlikely precisely because they require the final agent to receive such an implausibly strong private signal.

### Heterogeneous populations

The results above show that the magnitude of future-weighting has a substantial effect on the social interactions in homogeneous groups in which all agents share the same discount factor *γ*. However, real groups may be heterogeneous in their discounting of the future. For example, mixed species groups may be composed of animals with differing life cycles or metabolisms, who thus weight immediate and distant rewards differently. In single species groups, individuals of different ages may also attribute differing weights to future rewards, either because of differing expected future lifespans or because fitness may be disproportionately affected by success at specific ages. How will individuals in such heterogeneous groups interact so as to maximise their rewards over the timescales that are important to them?

Mixed populations may occur in many configurations, and deriving the optimal strategy for all individuals in such groups is analytically and computationally complex. Here I focus on a specific example to illustrate how differences in future-weighting affect the behaviour of rational agents. I consider a group drawn primarily from a homogeneous population sharing the same high value of *γ* = 0.99, representing agents with a typical time horizon of 100 future games. I assume that these agents, being drawn from a population of identical individuals, have adopted the optimal strategy for a homogeneous group with *γ* = 0.99. Within this group I place one further ‘invading’ agent with a short time horizon, *γ* = 0, representing an individual who prioritises immediate rewards. I assume that this agent is aware of the strategy employed by other group members, and derive its optimal response to this social environment. To avoid uncertainty, I continue to assume agents are otherwise identical, such that they share a common value of *ϵ* determining the quality of their private signals.

Figure 4 shows how the strategies of the two types of agents within this group differ. Each strategy is represented by the probability that an agent will choose option A, conditioned on the values of *n*_*A*_ and *n*_*B*_ (the number of previous agents choosing A and B respectively) and assuming that A is the true correct choice. In panel A these strategies are characterised as a function of *n*_*A*_ *− n*_*B*_. The red line indicates this probability for the agents with long time horizons (*γ* = 0.99); this shows very strong social feedback, such that for values of |*n*_*A*_ *− n*_*B*_| *≥* 2 agents follow the majority of previous decisions with probabilities very close to 1. The decision probability for the single agent with *γ* = 0 (who may be placed at any point in the sequence of decisions) is shown in blue. This agent shows a distinct pattern of social response: the agent is likely to follow the majority of previous decisions when this is in favour of A (albeit slightly less strongly than other agents); however, even when the majority is strongly in favour of B this agent remains as likely to choose A as B. This can be compared to the optimal strategy for a homogeneous group of agents with *γ* = 0, shown in black, where the response depends strongly on exactly how many agents prefer each choice, with a strong probability to choose B when this is favoured by a large majority of previous agents. This is because the agent with *γ* = 0 embedded in a group of agents with *γ* = 0.99 is aware of the strong social response of other individuals in the group. Since these other agents almost deterministically follow each other, the focal agent effectively views the majority choice as equivalent to a decision by one agent. The strong dependencies between the decisions of other agents limit how much information these can provide, and thus the agent who prioritises immediate rewards is forced to rely more on its private information, thus remaining open to environmental signals that contradict the behaviour of other agents. Panel B shows the strategy for the invading agent with *γ* = 0 respectively in more detail, as a functions of *n*_*A*_ and *n*_*B*_ (the equivalent visualisations for the homogeneous populations with *γ* = 0 and *γ* = 0.99 are shown in Figure 1A and C respectively). A notable feature is that the invading agent is most likely to choose option B (the incorrect choice) when a small majority of previous agents favour this option, rather than when all do. This agrees with the previous results shown in Figure 3: when other agents are known to prioritise consensus, disagreement in previous decisions is more convincing than consensus.

**Figure 4.**
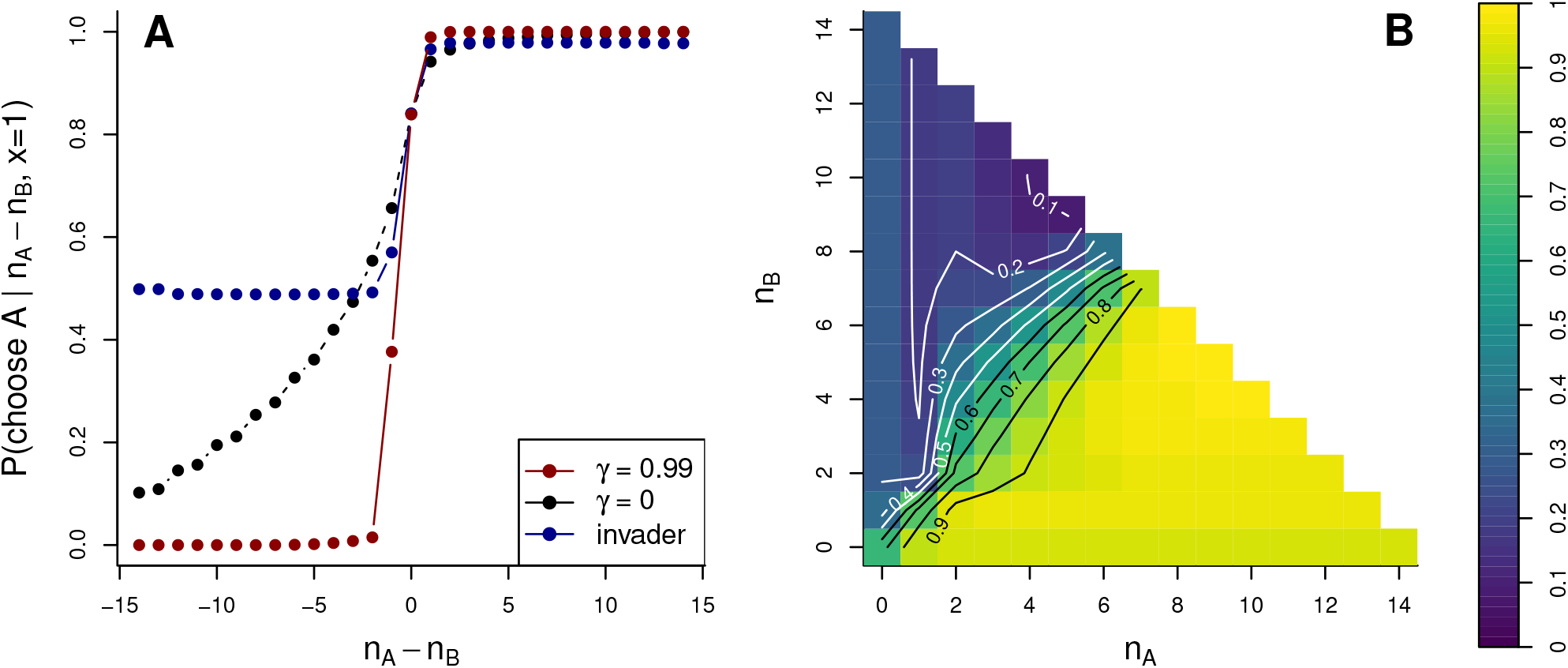
Optimal strategy in a mixed group. (A) Each strategy is characterised by the probability for an agent to choose the correct option A (*x* = 1), conditioned on the difference in the number of agents previously choosing options A and B (*n*_*A*_ *− n*_*B*_). The red line shows the optimal strategy for agents in a homogeneous group where all agents use future weighting *γ* = 0.99, implying a time horizon of 100 games. This strategy is characterised by an extremely strong social response, such that agents almost always follow the majority of previous decisions. The black line shows the optimal strategy for agents in a homogeneous group where all agents use future weighting *γ* = 0, implying no consideration of future rewards. Here agents follow others more weakly, but social responses are strong when the majority of previous decisions is large. The blue line indicates the optimal strategy for a single ‘invader’ agent with *γ* = 0 in an otherwise homogeneous group employing the optimal strategy for *γ* = 0.99. Under this strategy, agents attend only very weakly to the precise number of agents choosing each option. Notably, a single agent with *γ* = 0 remains highly likely to choose the correct option A even when there is a strong majority in favour of B. (B) The strategy for the invading agent with *γ* = 0 can be characterised in more detail by evaluating the probability that an agent chooses option A conditioned on the specific values of *n*_*A*_ and *n*_*B*_. Notably, the invading agent is most likely to choose B when a small majority of previous agents favour this option, in agreement with the results presented in Figure 3.

If one assumes that an agent’s value of *γ* is cryptic, the result above has an interesting corollary. As shown, most agents in the group will almost certainly follow a majority of previous decisions by other agents. However, the agent prioritising immediate rewards alone has a significant probability of disagreeing. Where the pre-existing majority is small, this implies that agents with shorter time horizons may change the eventual direction of a collective choice that otherwise would deterministically follow from whichever decision is made by the first agent.

## Discussion

Efforts to discern decision-making rules from first principles have predominantly concentrated on isolated decisions, overlooking the potential ramifications these decisions may have on the dynamics of group composition. The pivotal finding of this study demonstrates that where decisions occur not in isolation but as part of a continuous life history of the agent, and where these decisions determine later group composition, rational agents should display stronger sociality than would be expected in the case of isolated decisions. That is, where an agent will lose later benefits of group membership if it makes a decision at odds with its peers, it should be expected to more frequently follow what others have decided even at the cost of losing what it perceives to be immediately available rewards. This push towards stronger sociality emerges solely from considering the long-term decision-making success the agent can expect and is independent of any intrinsic social benefits [22] or risk reduction [23] the group can provide, and does not depend on mechanisms of collective conflict resolution that assume a consensus decision is required.

The results presented here provide an important context for considering the rationality of observed social behaviours. One example is the existence of despotic decision-making in animal groups with social hierarchies [24]. If a dominant individual leads a group to a food patch they can monopolise, why would lower-rank individuals follow them rather than foraging elsewhere? Previous work on groups of baboons has proposed that maintaining a social relationship with a dominant male may bring future benefits to other members of the troop, such as increased infant survival or protection from predators [24], which can outweigh the immediate loss from foraging. In addition to these benefits, the model presented here suggests that considering the outcome of a isolated foraging decision may be misleading. Although a single decision may present a clear choice between patches of obvious value, the baboons’ social responses should also favour maintaining group cohesion that can be beneficial in making future foraging decisions when identifying good foraging patches may be less straightforward. Another area where existing explanations of collective behaviour may be confounded by the theory presented here is in the concept of the Selfish Herd [23]: the idea that individuals seek to move towards the centre of a group to reduce risk of predation. Individual heterogeneity in risk-reward calculus has been posited as explaining the relative positions of individuals within groups, with more risk-tolerant individuals on the exterior of the group where they can take advantage of foraging opportunities [25, 26]. However, risk tolerance may be synonymous with shorter time horizons when considering future rewards, for instance in hungrier individuals; the model presented here suggests such individuals should be more willing risk leaving the group to obtain immediate rewards, which could result in them being found at the edge of the group independent of predation risk. This could also result in dynamics that resemble ‘leading according to need’ [27], as those individuals with the most pressing need of immediate rewards discount future rewards strongly and thus become apparently ‘socially indifferent’.

How much an agent can expect to benefit from continued group membership depends on how many future decisions that agent expects to face, and how strongly it discounts future versus immediate rewards. These two considerations are jointly represented by the future weighting *γ* in the model presented here; in reality they may both vary as a result of many factors. For example, animals that forage daily before regrouping at night may be assured of recovering group benefits within a relatively short time if lost during the day. Conversely, animals undergoing long-distance migrations may become isolated for very long periods or even permanently if they deviate from the main group. The results in this paper suggest that group cohesion should be greater in circumstances where group breakdown is irrevocable or very long term. As such, we might expect to see significant amounts of group breakdown in short term foraging (e.g fission-fusion collective behaviour [28, 29]), and more rigorous group cohesion in migration.

The magnitude of future-weighting can be expected to vary based on an individual’s life stage and life history. For instance, an animal that has not fed for a substantial time is likely to heavily discount the possibility of obtaining higher foraging rewards in future, preferring the higher chance of immediate food in the present. Life stage can also affect future reward discounting: empirical evidence points to a decreasing discount rate across the lifespan in humans [30]. How the weighting of future rewards should be determined is not modelled in this paper. However, in populations where individuals vary in their future discounting and the time horizons they consider, this paper has shown that individuals who discount the future more strongly should rationally attend less to the opinions and decisions of others, and be more suspicious in particular of consensus opinions which they can rationally consider to result from a high prioritisation of group cohesion on the part of agents with longer-term horizons. This accords with empirical studies on the effect of individual personality on collective behaviour showing that ‘bold’ individuals who take more risks [31] (suggesting a prioritisation of newly-available rewards over long-term security) are less inclined to follow the decisions of others in the group (e.g. [32, 33]). The inverse correlation between boldness and sociality has been interpreted as the result of more risk-tolerant individuals being more willing to forgo the safety of the group [34]; this study suggests that a further contribution may be that individuals with greater risk-tolerance are those who more-strongly discount the future, and who are therefore less inclined to forgo immediate rewards in favour of the greater expected payoffs in the future in a larger group.

Prioritising cohesion with the group necessarily compromises the agents’ accuracy in individual decisions. This should lower our expectations regarding the magnitude of collective wisdom groups can benefit from in contexts where disagreement bears the cost of exclusion from the group. As well as tempering expectations regarding the collective accuracy of, for example, collective animal navigation, this is also an important consideration when considering the collective judgements of human communities. On both small scales such as friendship groups and professional committees, up to huge online communities, individuals often explicitly or implicitly risk future exclusion from groups that are important to them when they disagree with others (see, for example, the effect of homophily and disagreement on ‘unfriending’ in social media [35, 36]). The results shown in this paper demonstrate that, where agents act in their rational long-term interests, this is likely to create a substantial barrier to realising the collective intelligence in these groups and instead create ‘groupthink’ dynamics where consensus is prized above accuracy [37]. In such communities we should especially be suspicious of unanimous opinions or decisions, as these are likely to result from information cascades [9] in which few agents contribute any meaningful information to the collective decision. Although they occur more rarely, we can conversely place more trust in decisions reached by relatively small majorities of agents, since the existence of disagreement precludes information cascades and points to the incorporation of genuinely strong private signals.

The model developed in this paper considers agents that choose sequentially in a random order, with each agent making a forced choice in its designated turn. This stands in the tradition of other theoretical models of sequential decision-making [38]. However, real-world decisions are rarely taken in a pre-defined sequence; instead agents such as humans and other animals may make decisions whenever they are sufficient confident to do so. An alternative model takes agents as continuously accumulating information until reaching some certainty threshold (e.g. refs. [39, 40]). A notable feature of this approach is that it predicts that early decision-makers will be those with better private information. Future research uniting these two modelling choices is likely to result in further insights into collective decision-making, and should consider the life-history perspective explored in this work. A further limitation of the model developed here is that all decisions both now and in the future are assumed to be equally difficult and of commensurate value. In practice an agent may be more willing to sacrifice rewards when faced with a relatively inconsequential decision now if that means having access to collective wisdom for more important decisions later. A detailed model of life history could incorporate decisions of differing types to investigate the effect this has on social conformity more broadly.

The core result of this paper is a reminder that while it may be either mathematically or empirically practical to study animal behaviours in isolation, the true consequences of behaviour play out over whole lifespans and in evolutionary time. The model in this paper remains reductive in that it concentrates only on the expected total abstract ‘reward’ that an agent can receive, and assumes that this maps straightforwardly on to fitness [41]. Such a translation should not be taken for granted: research on the economics of non-ergodic systems has demonstrated that what appear to be biases or irrational behaviours are often rational from the perspective of an agent’s life history [42]. In animal behaviour we should in particular expect that the translation of rewards such as food, safety and mating opportunities into fitness will be highly stochastic, non-linear and vary across a lifespan. Future attempts to identify rational behaviours from normative first principles should seek to make the link between behaviour and fitness more explicit and grounded in biological reality.

## Acknowledgments

This work was supported by UK Research and Innovation Future Leaders Fellowship MR/S032525/1 & MR/X036863/1 and the Templeton World Charity Foundation Inc. TWCF-2021-20647.

## References

[1] J. Krause and G. Ruxton, Living in Groups. Oxford University Press, Oxford, UK, 2002.

[2] D. J. T. Sumpter, Collective Animal Behavior. Princeton: Princeton University Press, 2010.

[3] I. D. Couzin, “Collective cognition in animal groups,” Trends in Cognitive Sciences, vol. 13, no. 1, pp. 36–43, 2009.

[4] T. Sasaki and S. C. Pratt, “Groups have a larger cognitive capacity than individuals,” Current Biology, vol. 22, no. 19, pp. R827–R829, 2012.

[5] A. Berdahl, C. J. Torney, C. C. Ioannou, J. J. Faria, and I. D. Couzin, “Emergent sensing of complex environments by mobile animal groups,” Science, vol. 339, no. 6119, pp. 574–576, 2013.

[6] S. Dall, L. Giraldeau, O. Olsson, J. McNamara, and D. Stephens, “Information and its use by animals in evolutionary ecology,” Trends in Ecology & Evolution, vol. 20, no. 4, pp. 187–193, 2005.

[7] T. J. Valone, “From eavesdropping on performance to copying the behavior of others: a review of public information use,” Behavioral ecology and sociobiology, vol. 62, pp. 1–14, 2007.

[8] V. Guttal and I. D. Couzin, “Social interactions, information use, and the evolution of collective migration,” Proceedings of the National Academy of Sciences, vol. 107, no. 37, pp. 16172–16177, 2010.

[9] S. Bikhchandani, D. Hirshleifer, and I. Welch, “A theory of fads, fashion, custom, and cultural change as informational cascades,” Journal of political Economy, vol. 100, no. 5, pp. 992–1026, 1992.

[10] A. Pérez-Escudero and G. G. De Polavieja, “Collective Animal Behavior from Bayesian Estimation and Probability Matching,” PLoS Computational Biology, vol. 7, no. 11, p. e1002282, 2011.

[11] S. Arganda, A. Pérez-Escudero, and G. G. De Polavieja, “A common rule for decision-making in animal collectives across species,” Proceedings of the National Academy of Sciences, vol. 109, pp. 20508–20513, 2012.

[12] R. P. Mann, “Collective decision making by rational individuals,” Proceedings of the National Academy of Sciences, vol. 115, no. 44, pp. E10387–E10396, 2018.

[13] R. P. Mann, “Collective decision-making by rational agents with differing preferences,” Proceedings of the National Academy of Sciences, vol. 117, no. 19, pp. 10388–10396, 2020.

[14] R. P. Mann, “Optimal use of simplified social information in sequential decision-making,” Journal of the Royal Society Interface, vol. 18, no. 179, p. 20210082, 2021.

[15] R. P. Mann, “Collective decision-making under changing social environments among agents adapted to sparse connectivity,” Collective Intelligence, vol. 1, no. 2, p. 26339137221121347, 2022.

[16] M. McPherson, L. Smith-Lovin, and J. M. Cook, “Birds of a feather: Homophily in social networks,” Annual review of sociology, vol. 27, no. 1, pp. 415–444, 2001.

[17] J. Figeac and G. Favre, “How behavioral homophily on social media influences the perception of tie-strengthening within young adults’ personal networks,” New Media & Society, p. 14614448211020691, 2021.

[18] R. J. Aumann, “Agreeing to disagree,” The Annals of Statistics, vol. 4, no. 6, pp. 1236–1239, 1976.

[19] R. P. Mann, “Optimising collective accuracy among rational individuals in sequential decision-making with competition,” Collective Intelligence, vol. 2, no. 2, p. 26339137231176481, 2023.

[20] D. J. T. Sumpter and S. C. Pratt, “Quorum responses and consensus decision making,” Philosophical Transactions of the Royal Society B: Biological Sciences, vol. 364, no. 1518, pp. 743–753, 2009.

[21] A. J. W. Ward, J. E. Herbert-Read, D. J. T. Sumpter, and J. Krause, “Fast and accurate decisions through collective vigilance in fish shoals,” Proceedings of the National Academy of Sciences, vol. 108, no. 6, pp. 2312–2315, 2011.

[22] L. Conradt and T. J. Roper, “Conflicts of interest and the evolution of decision sharing,” Philosophical Transactions of the Royal Society B: Biological Sciences, vol. 364, no. 1518, pp. 807–819, 2009.

[23] W. D. Hamilton, “Geometry for the selfish herd,” Journal of Theoretical Biology, vol. 31, no. 2, pp. 295–311, 1971.

[24] A. J. King, C. M. Douglas, E. Huchard, N. J. Isaac, and G. Cowlishaw, “Dominance and affiliation mediate despotism in a social primate,” Current Biology, vol. 18, no. 23, pp. 1833–1838, 2008.

[25] J. A. Teichroeb, “Vervet monkeys use paths consistent with context-specific spatial movement heuristics,” Ecology and Evolution, vol. 5, no. 20, pp. 4706–4716, 2015.

[26] M. del Mar Delgado, M. Miranda, S. J. Alvarez, E. Gurarie, W. F. Fagan, V. Penteriani, A. di Virgilio, and J. M. Morales, “The importance of individual variation in the dynamics of animal collective movements,” Philosophical Transactions of the Royal Society B: Biological Sciences, vol. 373, no. 1746, p. 20170008, 2018.

[27] L. Conradt, J. Krause, I. Couzin, and T. Roper, ““Leading According to Need” in Self-Organizing Groups.,” The American Naturalist, vol. 173, no. 3, pp. 304–312, 2009. PMID: 19199520.

[28] I. D. Couzin and M. E. Laidre, “Fission–fusion populations,” Current biology, vol. 19, no. 15, pp. R633–R635, 2009.

[29] C. Sueur, A. J. King, L. Conradt, G. Kerth, D. Lusseau, C. Mettke-Hofmann, C. M. Schaffner, L. Williams, D. Zinner, and F. Aureli, “Collective decision-making and fission–fusion dynamics: a conceptual framework,” Oikos, vol. 120, no. 11, pp. 1608–1617, 2011.

[30] L. Green, A. F. Fry, and J. Myerson, “Discounting of delayed rewards: A life-span comparison,” Psychological science, vol. 5, no. 1, pp. 33–36, 1994.

[31] D. Réale, S. M. Reader, D. Sol, P. T. McDougall, and N. J. Dingemanse, “Integrating animal temperament within ecology and evolution,” Biological reviews, vol. 82, no. 2, pp. 291–318, 2007.

[32] J. L. Harcourt, T. Z. Ang, G. Sweetman, R. A. Johnstone, and A. Manica, “Social feedback and the emergence of leaders and followers,” Current Biology, vol. 19, no. 3, pp. 248–252, 2009.

[33] L. M. Aplin, D. R. Farine, R. P. Mann, and B. C. Sheldon, “Individual-level personality influences social foraging and collective behaviour in wild birds,” Proceedings of the Royal Society B: Biological Sciences, vol. 281, no. 1789, p. 20141016, 2014.

[34] L. A. Gartland, J. A. Firth, K. L. Laskowski, R. Jeanson, and C. C. Ioannou, “Sociability as a personality trait in animals: methods, causes and consequences,” Biological Reviews, vol. 97, no. 2, pp. 802–816, 2022.

[35] H. Kwak, S. Moon, and W. Lee, “More of a receiver than a giver: why do people unfollow in twitter?,” in Proceedings of the International AAAI Conference on Web and Social Media, vol. 6, pp. 499–502, 2012.

[36] M. M. Skoric, Q. Zhu, and J.-H. T. Lin, “What predicts selective avoidance on social media? a study of political unfriending in hong kong and taiwan,” American Behavioral Scientist, vol. 62, no. 8, pp. 1097–1115, 2018.

[37] I. L. Janis, Groupthink. Houghton Mifflin Boston, 1983.

[38] S. Bikhchandani, D. Hirshleifer, O. Tamuz, and I. Welch, “Information cascades and social learning,” tech. rep., National Bureau of Economic Research, 2021.

[39] A. N. Tump, T. J. Pleskac, and R. H. Kurvers, “Wise or mad crowds? the cognitive mechanisms underlying information cascades,” Science Advances, vol. 6, no. 29, p. eabb0266, 2020.

[40] B. Karamched, M. Stickler, W. Ott, B. Lindner, Z. P. Kilpatrick, and K. Josić, “Heterogeneity improves speed and accuracy in social networks,” Physical Review Letters, vol. 125, no. 21, p. 218302, 2020.

[41] J. B. Silk, “The adaptive value of sociality in mammalian groups,” Philosophical Transactions of the Royal Society B: Biological Sciences, vol. 362, no. 1480, pp. 539–559, 2007.

[42] O. Peters, “The ergodicity problem in economics,” Nature Physics, vol. 15, no. 12, pp. 1216–1221, 2019.

